# Closed-loop acoustic stimulation modified by cathodal tDCS is beneficial for retention

**DOI:** 10.64898/2025.12.19.695567

**Authors:** T. Hoefer, T. Hösel, K. Hansen, M. Mölle, C. Peifer, M. Bazhenov, L. Marshall

## Abstract

We investigated how combining transcranial direct current stimulation (tDCS) with closed-loop acoustic stimulation (CLAS) during slow-wave sleep (SWS) affects memory consolidation and sleep-related neural oscillations. Cathodal tDCS was used to slightly reduce cortical excitability, thereby simulating a shifted brain state during CLAS and allowing us to compare the effects of standard CLAS to CLAS delivered under altered cortical conditions (CmodCLAS). Twenty-three participants (mean age 21.3 ± 2.6 years) completed two experimental nights: one with CLAS alone and one with CLAS combined with cathodal tDCS (CmodCLAS). Overnight retention on declarative memory tasks and morning learning performance were assessed. Memory outcomes revealed that CmodCLAS, but not CLAS alone, significantly improved overnight retention on the figural paired-associate task. In the EEG, CmodCLAS shifted the time-locked response to acoustic stimulation toward more negative potential values at the frontal region. CmodCLAS also prolonged slow oscillation (SO) duration at frontal sites while shortening SO duration at occipital sites, an effect not observed during standard CLAS. These findings demonstrate that the baseline level of cortical excitability during sleep modulates both the cognitive and electrophysiological effects of CLAS. They highlight the importance of brain state for non-invasive CLAS during sleep and suggest that CmodCLAS may serve as a useful approach for enhancing prefrontal cortical function during SWS.

**Significance statement:** Non-invasive brain stimulation (NIBS) is a promising approach to modulate brain activity and function. Unfortunately, NIBS studies are challenged by heterogenous findings. In addition to protocol differences, variability may depend strongly on the brain’s baseline activity (or ‘brain state’). Here, we used transcranial direct current stimulation (tDCS) to simulate a different brain state while applying closed-loop acoustic stimulation (CLAS) during slow-wave sleep (SWS). Sleep has been shown to enhance overnight memory retention. Results demonstrated significantly improved retention performance for cathodal tDCS modulated CLAS (CmodCLAS) as compared to CLAS alone. Furthermore, electrophysiological responses time-locked to the acoustic stimulus and spontaneous electrophysiological rhythms of SWS suggest that hippocampo-cortical function was specifically enhanced by CmodCLAS. Thus, underscoring the potential therapeutic relevance of CmodCLAS.

## Introduction

Interest in facilitating the brain’s electric activity during sleep has grown due to the wide range of functions attributed to sleep-related oscillations. Over the last half century, the potential role of sleep for memory consolidation has been intensely investigated (Rasch and Born, 2013; Klinzing et al., 2019; Brodt et al., 2023). Noninvasive brain stimulation (NIBS) studies have, particularly targeted rhythms and events associated with NREM sleep, both to probe their functional relevance as well as to investigate causality (Campos-Beltrán and Marshall, 2017; Grimaldi et al., 2020). During sleep, weak electric stimulation (constant or alternating current) and acoustic stimulation represent predominant NIBS approaches, supplemented more recently by targeted memory activation. Whereas the latter employs cues previously associated during wakefulness with information to be consolidated, closed-loop acoustic stimulation (CLAS) uses stimuli not associated with learning – analogous in this regard to weak transcranial electric stimulation (Malkani and Zee, 2020). The mechanisms through which these forms of electric and acoustic stimulation influence brain activity differ: weak electric stimulation primarily modulates cortical excitability, synaptic plasticity, and/or cortical-subcortical functional connectivity (Filmer et al., 2014; Bradley et al., 2022; Park et al., 2023), whereas slow oscillation (SO) CLAS appears to facilitate coupling across thalamo-cortico-hippocampal circuits (Ong et al., 2018; Aksamaz et al., 2024; Reith et al., 2025). Constant weak electric stimulation applied to the scalp (transcranial direct current stimulation, tDCS), modulates neuronal excitability and plasticity (Jackson et al., 2016; Yavari et al., 2018; Ren et al., 2022) potentially also through effects on neuromodulatory tone (Adelhöfer et al., 2019; Mishima et al., 2019).

A rather consistent observation across studies using weak electric stimulation, but also CLAS during sleep is that effects on electrophysiological brain activity are typically more robust and reliable than effects on behavior - such as memory consolidation or the homeostatic enhancement of post-sleep learning (Wiethoff et al., 2014; Dyke et al., 2016; Horvath et al., 2016; Papalambros et al., 2017; Henin et al., 2019; Ong et al., 2019; Papalambros et al., 2019; Malkani and Zee, 2020; Koo-Poeggel et al., 2022; Esfahani et al., 2023).

Given promising yet variable behavioral outcomes, it is essential - both for basic research and for clinical applications to understand the sources of variability in responsiveness to NIBS during sleep. Given the low risk involved, NIBS is particularly attractive as a possible means of treating mental health problems in young adults or even adolescents during sleep (Chellappa and Aeschbach, 2022; Meyer et al., 2024).

In addition to stimulation procedures and parameters, distinctions in the properties of neurophysiological targets, characteristics of the subject population, the brain state at the time of stimulation, and inherent inter-individual differences all appear to play significant roles in determining stimulation efficacy (Dedoncker et al., 2016; Dehnavi et al., 2021; Bradley et al., 2022; Navarrete et al., 2022; Alipour et al., 2024; Tabikh et al., 2025).

In the present experiment, we combined tDCS with CLAS during sleep to examine how changes in brain state alter the electrophysiological and behavioral responsiveness to CLAS. Our aim was to simulate within-subject differences in brain state and asses how differences modulate CLAS efficacy. To induce these state shifts, we applied cathodal tDCS over dorsolateral prefrontal cortices with anodal return electrodes at parietal sites.

## Materials and Methods

### Subjects

Twenty-three (17 females) participants, mean age (mean ± SD) 21.3 ± 2.6, ranging from 19 to 29 years, right-handed (as confirmed by the questionnaire of Annett handedness (Annett, 1970), non-smoking subjects with good knowledge of the German language completed the study. Exclusion criteria were mental and neurological pre-existing conditions, use of medication affecting the activation level or hormone balance, except for contraceptives, family history of epilepsy, metallic implants, shift work and prior participation in studies employing similar cognitive tasks. Twenty-one participants had been excluded after the adaptation night, mostly for not reaching the inclusion criteria of 40 min N3 sleep within the first half of the night, one subject dropped out due to a headache and another was excluded due to a sleep onset latency > 2 h. Twenty-seven participants finished all three nights, whereby data of 4 subjects were subsequently excluded due to technical recording issues or because the minimal number of 140 triggers corresponding to CLAS were not met (e.g. because of frequent awakenings). Informed written consent was obtained from all participants before participation. Subjects were to abstain from drinking caffeine, teein and alcohol on the days of the experiment. Participants were reimbursed for their participation. Approval of the local ethics committee of the University of Luebeck in accordance with the Declaration of Helsinki was given for the study.

### Procedure

After an adaptation night, the two experimental conditions were pseudo-randomized and counterbalanced across subjects. The adaptation night served also to control for adequate SO-detection, and volume of the acoustic stimulation in N3 sleep was assessed. Furthermore, values for a delta-theta ratio to detect NREM sleep, and the delay time for CLAS were calculated offline. Adaption and the first experimental session were separated by a minimum of 2 and a maximum of 7 days. For female participants without hormonal contraceptives the adaptation session was scheduled at the beginning of their luteal phase (between the 15^th^ and 21^st^ day of cycle) (Alonso et al., 2021). The two experimental nights were separated by at least 21 days (∼ 28 days for cycling female subjects). After electrode application subjects learned on two declarative (hippocampus-dependent) tasks in the experimental nights. Psychometric control parameters were assessed shortly before going to bed. Forty minutes after morning awakening, subjects performed the recall tasks and, in addition, learned on another hippocampus dependent task. Psychometric control tasks were performed, as indicated in Figure 1A.

**Figure 1.**
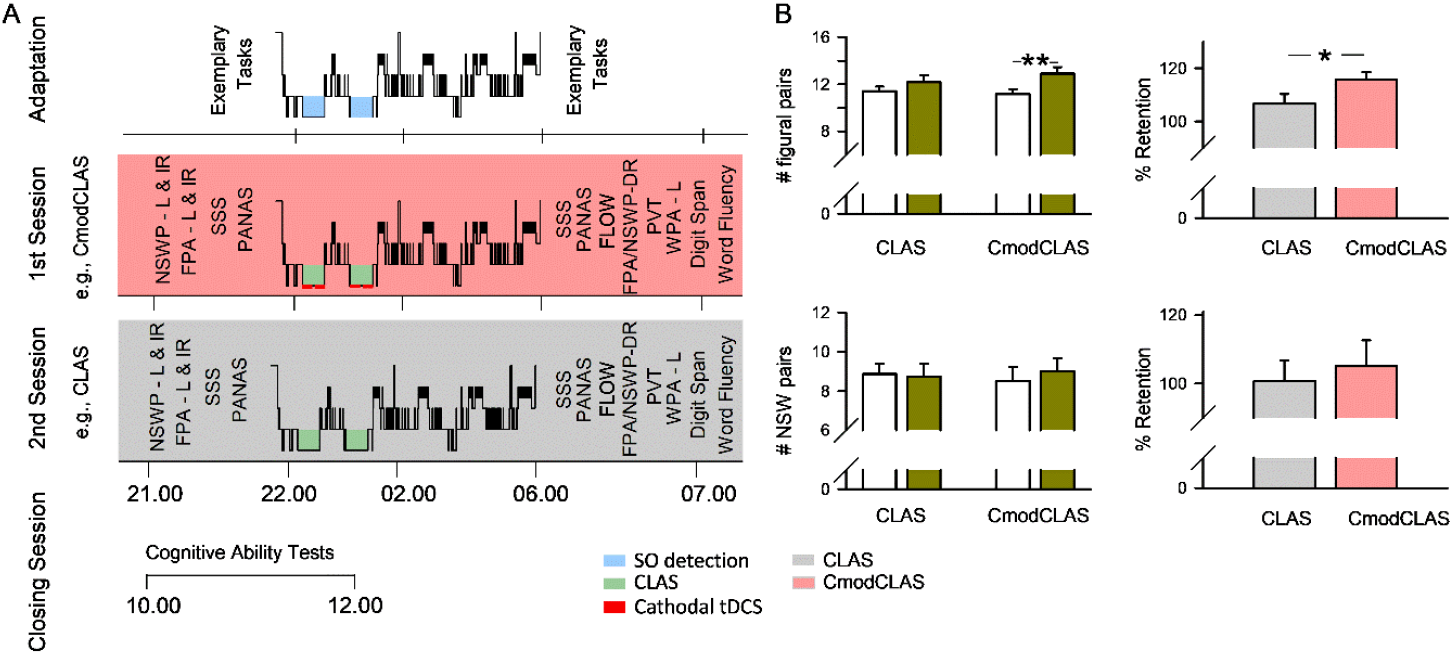
Experimental procedure and behavioral performance. **A** Time course and of the experimental procedure. **B.** Retention performance for the figural-paired task (N = 23) and nonsense word (NSW) paired task (N = 22). Left: Number of pairs at Learning (open bars) and recall (green filled bars) for CLAS and CmodCLAS. Right: Percent retention performance, with learning performance set to 100%. CmodCLAS: cathodal transcranial direct current stimulation (tDCS) modulated closed-loop acoustic stimulation (CLAS); DR: delayed recall; FPA: figural paired-associate task; IR: immediate recall; L: learning; NSWP: nonsense-word paired-associate; PANAS: positive and negative affect scale; PVT: psychomotor vigilance test; SO, slow oscillation; SSS: Stanford sleepiness scale; WPA: word pair-associate task. *p < 0.05, ** p < 0.01, Wilcoxon signed-rank test.

### Memory tasks and experiential state

Two declarative memory tasks (figural paired-associate, FPA and nonsense-word paired-associate, NSWP), and one task to assess encoding performance in the morning (word-paired-associate (WPA) were conducted. The FPA was previously described (Koo et al., 2018; Koo-Poeggel et al., 2022), and consisted of 16 pairs of figures (geometric and non-geometric lines and shapes) that were presented in two runs in a pseudorandom order for learning. Pairs were displayed for 4 s (interstimulus interval 500 ms). Immediate recall required selecting the match via mouse click from 7 other figures. There was no time limit, and during the second run 2-second feedback was given of the correct pair. The NSWP task consisted likewise of 16 non-word-pairs of two syllables each (e.g., GEMKIN, KRASUM), that were presented for 3 s in the identical fashion as the figural pairs. Immediate and delayed recall were conducted as in the FPA task, but with only one learning trial. Figures and nonsense words were adapted from (Sturm and Willmes, 1999). Retention performance was defined as the percentage of correctly recalled items at delayed recall minus the number of correct responses at immediate recall (mean of both immediate recalls for FPA) relative to learning performance.

Encoding performance in the morning was measured by a word-paired associate task (WPA), adapted from Koo et al. (2022). The test consisted of a total of 80 semantically unrelated pairs of German nouns, a cue and a target word presented for 3 s each (inter-stimulus interval of 500 ms). At immediate recall subjects responded to each cue word by speaking the target word the into a microphone, without time pressure. This procedure of encoding and immediate recall was repeated three times. No feedback was given. Learning performance on the WPA was defined as the percentage of the number of correct words recalled on each of the three trials relative to the total number of word pairs.

Before the above-described tests of memory recall and learning, we conducted a mental arithmetic task which contained 50 arithmetic equations per condition such as 3203 + 59 or 4285 – 14. After the completion of this task, we assessed participants’ flow experience during the task, defined as the enjoyable state of being fully immersed in an optimally challenging task (Peifer and Engeser, 2021). The 10 items of the Flow Short Scale (FSS; Rheinberg et al., 2003) used a 7-point used on questions covering two components of flow experience: absorption and fluency, that together reflect overall flow experience.

### Psychometric control data and cognitive ability

Before the lights out in the evening and before cognitive testing in the morning subjects completed the positive and negative affect scale (PANAS) (Watson et al., 1988) and Stanford sleepiness scale (SSS) (Hoddes et al., 1973). Furthermore, in the morning subjects conducted a 5-minute psychomotor vigilance task (PVT) (Roach et al., 2006), the Regensburger verbal fluency test (RWT) (Aschenbrenner and Lange, 2001), Digit Span Forward (DSF) and Backward (DSB) test (Wechsler Adult Intelligence Scale (Wechsler, 1997). Tests to assess general fluid intelligence (APM, Ravens’s Advanced Progressive Matrices, (Raven et al., 2010)) and memory quotient score (MQ, (Bäumler, 1974)) took place in a subsequent morning session.

### Data acquisition

Electroencephalographic activity (EEG) was amplified and recorded from 13 sites according to the international 10-20 system (F7, F8, Fz, C3, Cz, C4, P3, Pz, P4, O1, O2, M1 and M2) together with ventral and horizontal electrooculogram (VEOG, HEOG) between 0.032 and 150 Hz (50-Hz notch filter; LiveAmp, Brain Products GmbH, Gilching, Germany). Cz served as reference, and Afz as ground. EEG data were re-referenced to linked mastoids for analyses. EMG was amplified between 5.305 and 100 Hz. Electrocardiography (ECG; right arm to anterior axillary line at the level of the fifth intercostal space (V5), was amplified between 1 Hz and 25 Hz. Impedance was kept below 5 kOhm for EEG, HEOG, VEOG and below 10 kOhm for EMG and ECG. All data were sampled at 500 Hz.

### Slow oscillation (SO) detection and closed-loop acoustic stimulation (CLAS)

CLAS targeted the SO Up-state during deep NREM sleep as previously described (Ngo et al., 2013; Koo-Poeggel et al., 2022; Esfahani et al., 2023; Reith et al., 2025). In brief, Fpz referenced to linked mastoids detected SOs online, with the EEG signal filtered between 0.25 Hz and 4 Hz (Digitimer, Hertfordshire, UK) at a sampling rate of 200 Hz. The delta-theta ratio was continuously calculated to determine NREM sleep but filtered between 0.25 Hz and 45 Hz.

CLAS was triggered when: (1) The EEG signal exceeded an adaptive amplitude threshold in negativity (updated every 0.5 s) based on the EEG amplitude of the preceding 5s, and initially set at - 80 μV. (2) The delta/theta ratio was above the individual threshold for NREM sleep. Our CLAS condition was identical to Hausdorf and colleagues (Hausdorf et al., 2025): CLAS was manually activated by the investigator after at least 4 min of stable N3 sleep for the first deep sleep cycle, and after 2 min of N3 for any subsequent sleep cycle. In the condition with CLAS modified by cathodal tDCS (CmodCLAS) electrical stimulation was initiated prior to CLAS, and acoustic stimuli (pink 1/f noise of 50 ms duration, including 5-ms of each rising and falling times) were only released after complete attenuation of electric stimulation artefacts (∼ 30 s). Similarly, CLAS was terminated shortly before the end of tDCS (approximately 30 s) to avoid artefacts in the event-related responses. In the CLAS condition acoustic stimulation was initiated after ∼4.5 min (4 min + ∼ 30 s) of stable N3 for the first sleep cycle and 2.5 min (2 min + 30 s) for any subsequent sleep cycle. Thus, initiating CLAS in both sessions at closely comparable periods (Hausdorf et al., 2025).

Acoustic stimulation consisted of two bursts of pink 1/f noise of 50 ms duration, including 5-ms of each rising and falling times delivered bilaterally through in-ear phones (MDR-EX15LP, Sony Germany). The second stimulus arrived 1075 ms after the first, that was delivered shortly before the SO positive peak. The minimal interstimulus interval between acoustic stimulation pairs was 2.5 s. Volume was initially set to 35 dBA, and was increased with sleep depth up to about 49 dBA. When the subject showed signs of arousal or shifted toward lighter sleep, CLAS was terminated, and the target volume was readjusted. Stimulation was limited to the first four hours of time in bed. We aimed for the same number of acoustic stimuli in the second session as had been applied in the first experimental session.

### Transcranial direct current stimulation

Cathodal tDCS was applied using two parallel circuits to sites F3 and F4 (each 9cm^2^, Neuroconn, DC-Stimulator Plus, Ilmenau Germany) with ipsilateral return electrodes (each 14 cm^2^) positioned between P3 and O1, and P4 and O2, respectively, and attached using conductive paste (Ten20 Conductive, Weaver and company, Aurora, USA). A ramp up and down of 8 seconds with a maximum current intensity of 1000 µA per circuit was used. Impedance was kept below 2 kOhm. TDCS was to be applied twice for 15 min, with a 5 min break within the first half of the night (first 4h of sleep), beginning after 4 min of stable N3 sleep of the first deep sleep cycle, and 2 min of stable N3 sleep of any subsequent sleep cycle. A stable delta-theta ratio above 20 at Fpz (short pauses of maximum 5 s were accepted) was required. An interval of tDCS was terminated if subjects switched to a lighter sleep stage. TDCS or sham-tDCS during N3 continued throughout the night until subjects had received 30 minutes in total. In sham-tDCS (CLAS only) 30-second epochs of tDCS were applied.

### EEG analyses

Polysomnography was conducted with SleepPilot (https://github.com/xuser/SleepPilot, v0.9.4-beta) by two independent researchers blinded for the condition, and used 30-s epochs of EEG-recordings (at Fz, C3, C4, Pz; bipolar VEOG, HEOG and EMG). Wake, N1, N2, N3 and REM sleep were determined according to American Academy of Sleep Medicine guidelines (Berry et al., 2012). In addition, movement arousals and artefacts were marked. Time frames of tDCS were marked for EEG analyses.

Data analyses were performed using Spike2 (Cambridge Electronic Design, Cambridge, England). To assess event-related responses (wide-band, slow and fast spindle activity) EEG signals were first low pass-filtered at 35 Hz (FIR filter, –3 dB at 32.0 Hz) and then downsampled to 100 Hz. Windows of -1 to 3 s time-locked to the first stimulus were extracted from artefact-free epochs NREM sleep periods (N2 and N3) and averaged. For spindle activity, before extracting the 4-s windows a FIR bandpass filter with a width of 3 Hz around individually determined spindle peak frequencies was applied (within 9 - 12 Hz for slow spindle activity; within 12 – 16 Hz for fast spindle activity). The root mean square (RMS) was calculated for each sample point and smoothed (time window: 0.2 s).

For offline SO identification, signals were first bandpass filtered (0.5 – 3.5 Hz). Negative and positive peak potentials were then detected from all intervals with zero-crossings separated by 0.75 to 2 s. Mean amplitude thresholds were subsequently determined for all negative and negative-to-positive peak potentials, from thresholds set at 1.25 times the mean negative or negative-to-positive peak potential, respectively. A SO was defined when for successive negative and negative-to-positive peak potentials amplitudes surpassed the respective thresholds. Durations of positive and negative SO half-waves were calculated using the zero-crossings of detected SOs.

Discrete spindles were detected from the low-pass filtered and downsampled EEG data. Spindle detection was adopted from Mölle et al. (2011). Individual fast (12 - 16 Hz) and slow (9 - 12 Hz) spindle peak frequencies were identified from power spectra at all channels for all artefact-free NREM sleep periods. Mean slow and fast spindle peak frequencies were 11.6 ± 0.7 Hz and 13.5 ± 0.5 Hz, respectively. After band-pass filtering (centered ± 1.5 Hz around detected individual peak frequencies) a RMS representation of the filtered signal was calculated using a sliding window of 0.2 s and a step size of one sample. Finally, RMS data were smoothed with a 0.2-s-size sliding window. Time frames were considered as spindle intervals if the RMS signal exceeded a threshold of 1.5 SD of the bandpass filtered signal for 0.5 - 3 s, and at least one sample point was above a threshold of 2.5 SD of the bandpass filtered signal. Detected spindles were merged if the intervening interval was < 0.5 s and the total RMS superseding the 1.5 SD threshold was < 3 s. Spindle count, density (i.e., number of spindles per 30 s), mean peak to peak amplitude, mean length and mean spindle RMS throughout NREM sleep were analyzed.

EEG power during NREM sleep was analyzed within the following frequency bands SO (0.488-1.4648 Hz), SWA (0.488 – 4.0283 Hz, delta (1.5869 - 4.0283 Hz) theta (5.004-8.911 Hz), slow spindle (10-12.5 Hz) and fast spindle (12.5-15 Hz)

### Statistical Analyses

The parameters of detected oscillatory events (SOs and spindles) were tested statistically by repeated measures analyses of variance (rmANOVA) with the factors condition, COND (levels: CLAS and CmodCLAS), and corresponding scalp topography, TOPO (levels: F7, Fz, F8, C3, Cz, C4, P3, Pz, P4, O1 and O2). In rmANOVA with 4 topographies, data were averaged across corresponding lateral and midline locations. Before statistical analyses individual data of time-locked SO and spindles RMS responses to acoustic stimulation were first baseline normalized (initial 1.5 seconds).

Time series analyses (time-locked SO, time-locked spindle RMS, and offline detected SOs) for comparisons between conditions were conducted by Wilcoxon signed-rank tests, with false positives controlled for by the Maris–Oostenveld permutation test. The latter incorporated all time points and 11 electrodes. Differences between conditions in SO-spindle event-event coupling were tested via Wilcoxon signed rank tests using the Benjamini-Hochberg procedure to control for false discovery rate (FDR).

Memory retention was tested by rmANOVA with the factors condition, COND (levels: CLAS and CmodCLAS), and time, TIME (levels: evening, morning). Subsequent to analyses of variance, differences in retention performance (morning delayed recall – evening immediate recall) between CLAS and CmodCLAS were assessed by paired sample tests without multiple comparison correction since this was a primary behavioral hypothesis. For the learning task a one-way rmANOVA with the factor Learning TRIALS (L1, L2, L3) was employed.

In general, for normally distributed data, as assessed by the Shapiro-Wilk test together with skewness and kurtosis, repeated measures variance analyses with subsequent paired sample t-Tests were performed. Non-parametric paired sample tests were employed for not normally distributed data. ANOVA results are Greenhouse-Geisser corrected to compensate for any violation of sphericity (uncorrected degrees of freedom are given). Bands of EEG power were logarithmized prior to statistical analyses to obtain normality.

Values are reported in mean ± SEM. A value p < 0.05 was considered significant. Statistical analyses were conducted with SPSS (IBM SPSS Statistic, version 22.0, IBM, Armonk, NY, USA) or R/RStudio (version 2025 4.5.1).

## Results

### CmodCLAS and behavior

The rmANOVA for the figural paired-associate (FPA) task revealed a significant COND × TIME interaction (F(1,22) = 5.503, p = 0.038; η^2^ = 0.182). There was also a main effect of TIME (F(1,22) = 18.881, p < 0.001; η^2^ = 0.462). However, overnight retention improved significantly only after CmodCLAS (T(22) = –5.428, p < 0.001), whereas CLAS alone showed only a trend toward improvement (T(22) = –1.897, p = 0.071). Percent retention was significantly higher with cathodal tDCS relative to CLAS alone (115.8 ± 2.7 vs. 106.8 ± 3.6; T(22) = –2.148, p = 0.043; Figure 1B). FPA learning performance in the evening did not differ between conditions (T(22) = 0.561, p > 0.5).

Performance on the NSWP task failed to show overnight improvement (F(1,21) = 0.070, p > 0.794 for main effect of TIME) nor any difference in overnight performance between tasks (F(1,21) = 1.228, p > 0.2, for COND x TIME). One subject’s NSWP data were incomplete and excluded.

Learning performance on the WPA task after morning awakening improved significantly across the three trials (F(2,44) = 275.3, p < 0.001), with MQ scores correlating with learning differences from the 1st to 2nd trial (r = 0.588, p = 0.003) and across all three trials (r = 0.535, p = 0.009), but not with the change from the 2nd to 3rd trial (p > 0.6). There was no main effect of COND and no COND × TIME interaction (p > 0.5).

The flow subscale fluency was significantly affected by CmodCLAS (T(21) = -2.618, p = 0.009) and also overall flow T(21) = -2.173, p = 0.03), but absorption was not affected (p > 0.5). CmodCLAS decreased both fluency and overall flow performance (fluency: 5.4 ± 0.2 vs. 4.9 ± 0.3).

### Modulation of EEG activity by CmodCLAS

Polysomnographic variables did not differ between conditions (Supplementary Table S1). Both, responses time-locked to acoustic stimulation and spontaneous EEG activity during offline detected SOs across entire NREM sleep showed sustained potential shifts under CmodCLAS relative to CLAS. The main electrophysiological of CmodCLAS on responses time-locked to the first acoustic stimulus was a sustained negative potential shift at the midline frontal site.

This negative shift commenced with the detected negative half-wave and persisted for ∼1.5 s, spanning the subsequent positive (Up) state, the interim negative (Down) state, and the following positive (Up) state elicited by the second acoustic stimulus. During acute cathodal tDCS, mean amplitudes of this shift appeared more pronounced. After permutation testing, significant effects were found only at Fz, both during acute tDCS (Figure2) and across all NREM N2 and N3 epochs (Supplementary Figure 1). Fast and slow spindle RMS responses were unaffected (Figure 3; Supplementary Figures 2, 3). The average number of NREM sleep epochs receiving acoustic stimulation did not differ significantly between CLAS (297.5 ± 23.9) and CmodCLAS (261.9 ± 17.0; p > 0.1).

**Figure 2.**
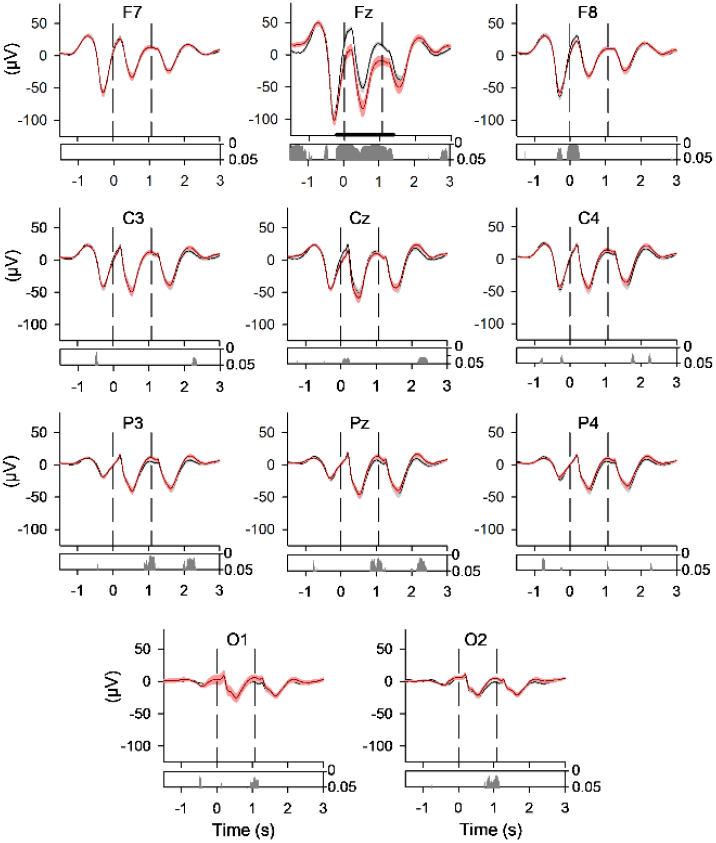
Stimulus-locked EEG responses to acoustic stimuli in CLAS and CmodCLAS for tDCS On time periods. Top diagrams: Grand mean waveforms (± SEM) at locations F7, Fz, F8, C3, Cz, C4, P3, Pz and P4 for CLAS (black) and CmodCLAS (red); baseline normalized data. Bottom diagrams: horizontal bars indicate time points of significant differences without multiple comparison correction (Wilcoxon signed-rank test); horizontal bars indicate significant clusters after permutation testing, p < 0.05). N = 23? Baseline normalization occurred over - 0.99 to -0.01 sec.

**Figure 3.**
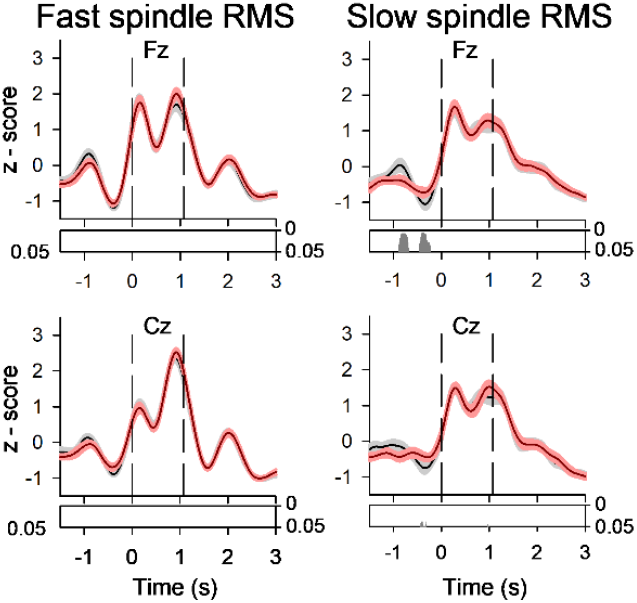
Stimulus-locked fast and slow spindle root mean square (RMS) to acoustic stimuli in CLAS and CmodCLAS. Top diagrams: Grand mean waveforms (± SEM) of stimulus-locked responses at the midline electrode Fz for CLAS (black) and CmodCLAS (red). Bottom diagrams: horizontal bars indicate time points of significant differences without multiple comparison correction (Wilcoxon signed-rank test, p < 0.05), N =23.

Offline-detected SOs across the entire NREM period, revealed time-dependent shifts in opposite directions. At Fz, three significant clusters were identified: In two, CmodCLAS shifted potentials toward more negative values relative to CLAS during the Up-state into the Up-to-Down state transition (p = 0.005) and the Down-to-Up transition into the next Up-state (p = 0.019). In the third cluster (p = 0.005), the subsequent negative-going wave showed higher positive potentials under CmodCLAS. Effects at F8 were in the same direction (uncorrected) but did not reach significance (Figure 4).

**Figure 4.**
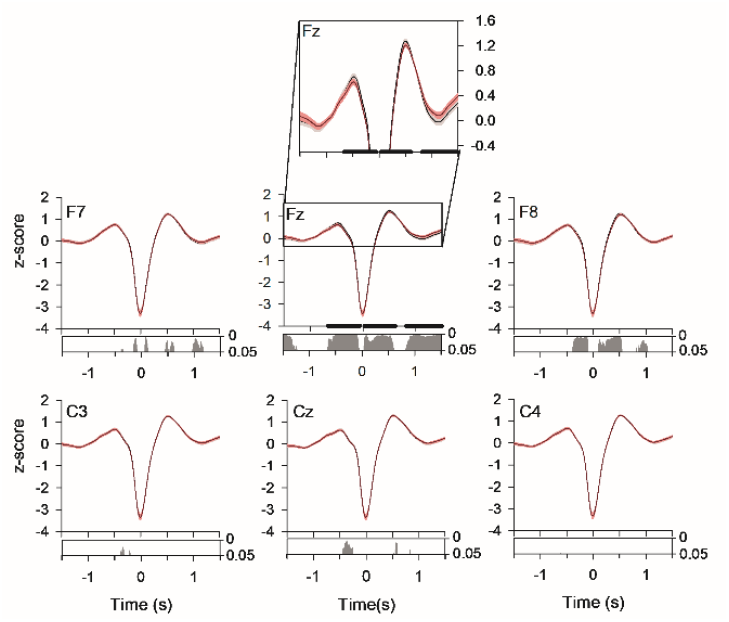
For all offline detected slow oscillations (SO) throughout NREM sleep, the effect on closed-loop acoustic stimulation (CLAS) of cathodal tDCS (CmodCLAS). Grand mean waveforms (± SEM) of spontaneous SO at locations F7, Fz, F8, C3, Cz, and C4 for CLAS (black) and CmodCLAS (red). Bottom diagrams: Horizontal bars indicate time points of significant differences without multiple comparison correction (Wilcoxon signed-rank test). *N* = 23.

CmodCLAS also significantly altered parameters of offline-detected SO events. SO length increased at frontal sites Fz and F8 (F(1,22) = 14.801, p ≤ 0.001) but decreased at occipital sites (F(1,22) = 7.046, p ≤ 0.014; overall COND × TOPO: F(10,220) = 9.353, p < 0.001; overall TOPO: F(10,220) = 16.728, p < 0.001; Figure 5). To examine these effects more closely, we compared positive and negative SO half-waves. At the two frontal locations, negative half-waves were significantly longer in CmodCLAS (T(22) ≥ |2.443|, p ≤ 0.023; COND: F(1,22) = 17.859, p < 0.001); with no significant distinction between effects at Fz and F8 (p > 0.16). At occipital sites, positive half-waves were significantly shorter (T(22) > 2.927, p ≤ 0.004; COND: F(1,22) = 10.500, p = 0.004; Figure 5).

**Figure 5.**
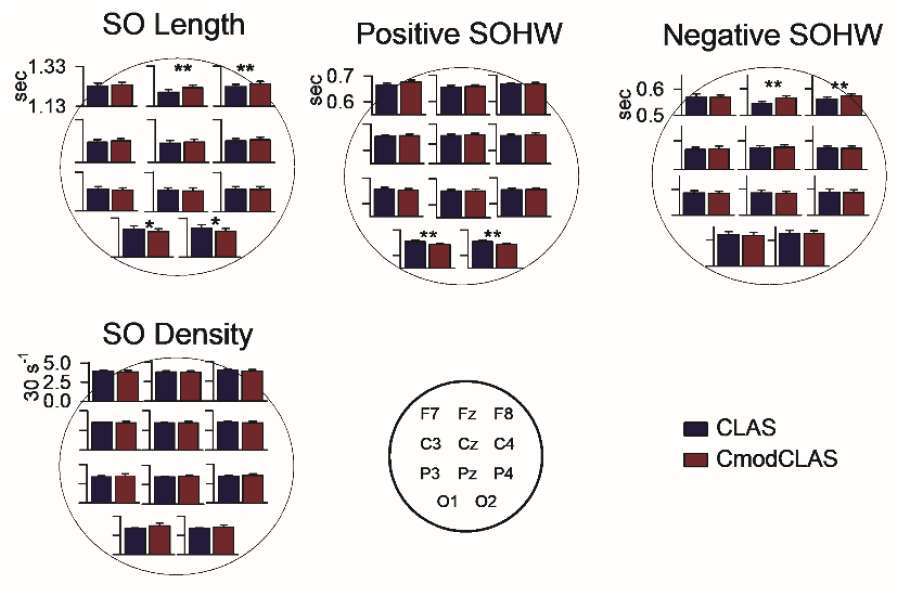
Effects on discrete SO measures of closed-loop acoustic stimulation (CLAS) and CmodCLAS. Measures revealing significant condition differences for discrete SOs in CLAS (blue bars) and CmodCLAS (red bars) throughout NREM sleep are shown. \**p* < 0.05, \*\**p* < 0.01 for post hoc tests; see text for rmANOVA results. *N* = 23.

A significant COND × TOPO interaction for SO density (F(10,220) = 2.686, p = 0.04) suggested opposite anterior–posterior effects, however paired comparisons were non-significant (T(22) < |0.634|, p > 0.5; Figure 5). CmodCLAS significantly increased the positive SO peak potential at occipital locations only (T(22) = |2.417|, p = 0.024; F(1,22) = 5.884, p = 0.024; COND × TOPO: F(10,220) = 2.742, p = 0.026). Neither the negative-to-positive SO slope (Down-to-Up transition) nor negative peak potentials were significantly affected (p > 0.07).

CmodCLAS did not influence any discrete slow or fast spindle metrics (p > 0.06). Supplementary Table S2 provides spindle density, duration, and RMS collapsed across conditions. EEG frequency-band power during NREM sleep was also unaffected (Supplementary Figure 4).

### Psychometric control parameters

As expected, PANAS revealed a significant main effect of Time, with higher positive affect in the morning than evening (2.8 ± 0.1 vs. 2.2 ± 0.1; χ^2^(1) = 30.425, p < 0.001). Positive affect was also higher under CmodCLAS (2.6 ± 0.1 vs. 2.4 ± 0.01; χ^2^(1) = 7.787, p = 0.005). Negative affect was not significantly influenced by Condition or Time. SSS scores showed a significant main effect of Time, with subjects less sleepy in the morning (2.6 ± 0.2 vs. 4.4 ± 0.2; χ^2^(1) = 37.866, p < 0.001; Supplementary Table S3). CmodCLAS did not affect SSS (p > 0.3, for COND main effect and COND x TIME interaction). Morning digit span and word fluency performance did not differ between conditions (p > 0.2).

## Discussion

This study examined how modest changes in cortical excitability and activity influence the brain’s responsiveness to CLAS during SWS. Inter-individual variability in responsiveness to CLAS may arise, in part, from differences in baseline cortical excitability. The ability to systematically alter susceptibility to CLAS through tDCS strongly supports this view. Numerous studies have underscored strong inter-individual differences in sleep rhythms, their coupling as well as differential susceptibility to non-invasive brain stimulation (Li et al., 2015; Cox et al., 2018; Koo et al., 2018; Kasten et al., 2019; Dehnavi et al., 2021, 2023; Lewis et al., 2025).

Using the same experimental design as in the present study, across all subjects, anodal tDCS failed to modify the behavioral response to CLAS (Hausdorf et al., 2025). Only subjects with high fluid intelligence scores responded to anodal tDCS modified CLAS as compared to CLAS alone, revealing impaired memory consolidation. In contrast, CmodCLAS improved consolidation across all participants on the figural paired-associate task. In this regard the modulation by tDCS of opposite polarity led to opposing effects on behavior, as expected. The improvement in consolidation with cathodal stimulation over the dorsolateral prefrontal cortex may seem counterintuitive, since cathodal stimulation is often associated with reduced cortical excitability and activity when applied over the motor cortex (Nitsche and Paulus, 2000), but see also Klees-Themens and Théoret (2023). In cognitive domains, evidence for the anodal-excitation and cathodal-inhibition pattern (AeCi) is far more heterogenous, with cathodal tDCS often increasing performance (Jacobson et al., 2012; reviewed in Brückner and Kammer, 2017; Schroeder and Plewnia, 2017). We suggest that CmodCLAS may have reduced neuronal background noise and/or interference under the cathodes (DLPFC), and at the same time enabled more efficient interregional communication on delivery of CLAS. Ultimately memory consolidation was enhanced.

The tendency for AmodCLAS and CmodCLAS to pose opposite changes in performance are less clearly reflected in electrophysiology. CmodCLAS shifted activity time-locked to the acoustic stimulus toward more negative values, and produced similar polarity shifts in offline detected SOs within ± 0.6 s of the negative half-wave peak. Approximately 1 s after this negative half-wave peak, offline detected SOs under CmodCLAS showed a shift toward more positive values. These effects were significant at the midline frontal electrode, with tendencies in the same directions occurring at the right frontal electrode. CmodCLAS also increased SO length at frontal sites. Interestingly, the negative polarity shifts and the increase in SO length at frontal sites were also observed with AmodCLAS. However, some subtle, yet likely crucial differences emerged. Firstly, following the time-locked acoustic stimulation CmodCLAS generated a stronger shift toward negative values compared to CLAS than AmodCLAS. Secondly, for offline detected SOs the negative shifts induced by CmodCLAS appeared topographically more restricted. Thirdly, in addition to increasing SO length at frontal electrodes, CmodCLAS led also to a pronounced decrease in SO length at occipital sites. In contrast, AmodCLAS increased SO length (for COND: F(1,19) = 7.744, p = 0.012; Hausdorf et al., 2025) without any significant topographical interaction (F(10,190 = 1.921, p = 0.113; for all NREM sleep epochs; result unpublished).

In our prior work (Hausdorf et al., 2025) we argued that the stronger modulation of spontaneous SO activity relative to time-locked acoustic responses implied that the specific sensory pathway was not primarily affected by AmodCLAS. Instead, we proposed that AmodCLAS modulated spontaneous “background” activity, potentially via an increase in neuromodulatory noradrenergic tone. Here, we speculate that the relatively stronger time-locked response induced by CmodCLAS, may reflect on the one hand a dampening of background cortical activity, and on the other hand, strengthened hippocampal input to the cortex. Short neutral sounds during sleep generate short (∼27 ms) and longer latency (hundreds of milliseconds) responses of hippocampal neurons (Vinnik et al., 2012; Rothschild et al., 2017). A recent, source-localized magnetoencephalography study suggests that CLAS engages non-lemniscal auditory pathways, with one model requiring low noradrenergic tone for the emergence of the N500-P900 component (Jourde et al., 2024). During active locomotion, acoustic stimuli were shown to reach the hippocampus via a non-canonical route, i.e., not relying on thalamocortical pathways (Winne et al., 2025). Whether and how such hippocampus responses to neutral sounds directly contribute to memory consolidation is still unclear.

Interestingly, flow experience, in particular the subscale fluency, was decreased upon awakening after CmodCLAS. Although at first glance this appears to contradict the beneficial effect of CmodCLAS on memory consolidation, the two outcomes may reflect the same underlying modulation of prefrontal dynamics. Flow experience while conducting a similar arithmetic task was associated with deactivation of medial prefrontocortical activity compared to boredom as assessed by regional cerebral blood flow (Ulrich et al., 2014; Harris et al., 2017; Ulrich et al., 2018). Thus, if the effect on memory consolidation relied on an increase in the signal-to-noise ratio, flow performance likely dependent on broader prefrontal engagement could be impaired. The failure of tDCS modulated CLAS in this and a previous study to affect morning learning on the WPA task may be technical, since only three learning trials were used (Hausdorf et al., 2025). In a previous study differential learning tended to emerge at later learning trial (Koo-Poeggel et al., 2022).

Taken together, so far, we propose that CmodCLAS induced dampening of prefrontal cortical activity during sleep may facilitate task-relevant hippocampal-cortical communication by improving the signal-to-noise ratio of either incoming hippocampal information or by enhancing task-relevant cortico-hippocampal functional interactions.

A further major difference from AmodCLAS was the opposite effect of CmodCLAS on SO length between frontal and occipital regions: SO length increased at frontal sites but was significantly reduced at occipital sites near the return (anodal) electrodes. No other topographical differences between conditions in positive or negative SO half-wave length were found. This is notable, as the active anodal electrode in AmodCLAS increased SO length globally. Ignoring the different electrode sizes of the active and return electrodes, applied current intensity and electric field properties, our results nonetheless show opposite effects on SO length in response to stimulation of the same nominal polarity. Whether increased posterior positive SO halfwave length (Up-state) specifically aided consolidation of the figural (as opposed to the verbal) paired associate task warrants further investigation. A more holistic view would be to attribute the differential response at the occipital region to CmodCLAS and AmodCLAS exerting contrasting influences on anterior-posterior cortical functional connectivity.

In summary, CmodCLAS used here to simulate within-subject shifts in brain state, enhanced consolidation of a figural paired-associate task, shifted the EEG toward more negative values at frontal sites, increased SO duration at fronto-central sites, but reduced SO duration at occipital locations. Neither spindle activity nor SO-spindle coupling were altered. Given that acoustic stimuli and CLAS during sleep induce hippocampal responses, our findings suggest the beneficial effects of CmodCLAS on memory consolidation may arise from improved (bidirectional) hippocampal-prefrontocortical communication (Helfrich et al., 2019; Sanda et al., 2020).

If confirmed by future studies, the combination during SWS of cathodal tDCS and CLAS suggesting a shift in the balance of excitatory-inhibitory cortical network activity together with increased hippocampo-cortical communication could indeed be of high clinical relevance. Functional changes in cortical excitability, impaired hippocampal function and hippocampo-cortical communication are implicated in many mental disorders such as ADHD, anxiety and depression. These conditions involve changes in neural plasticity, frequently co-occur, and urgently need effective treatments – particularly non-pharmacological interventions suitable for younger age groups (Bramson et al., 2023; Kuo et al., 2024; Fu et al., 2025; Woodham et al., 2025).

## Acknowledgements

We thank Eric Gromodka, Shu Zhang and Dr. Ping Koo-Poeggel for graphical and technical assistance. This work was supported by the US-German Collaboration in Computational Neuroscience (NSF/BMBF grants 01GQ1706 and IIS-1724405), the German Research Society, DFG (MA 2053/9-1; 2053/11-1; MA 2053/12-1), National Science Foundation (grant EFRI BRAID: 2223839), and National Institutes of Health (grants 1RO1MH125557, 1R01NS109553, RF1NS132913).

Data is accessible upon reasonable request.

The authors have no conflict of interest to declare.

## Notes

### Competing Interest Statement

The authors have declared no competing interest.

