## Supplementary Information for "Closed-loop acoustic stimulation modified by cathodal tDCS is beneficial for retention"

### Supplementary Table S1 Sleep parameters

| | CLAS | CmodCLAS |
|---|---|---|
| TST (min) | 437.1±8.2 | 437.0±6.7 |
| WASO (%) | 0.0±0.0 | 0.0±0.0 |
| N1 (%) | 6.0±1.3 | 5.2±0.7 |
| N2 (%) | 49.9±1.3 | 51.0±1.5 |
| N3 (%) | 26.3±1.6 | 26.1±1.9 |
| REMS (%) | 17.7±1.0 | 17.8±1.0 |
| Sleep onset latency (min) | 16.2±2.0 | 13.7±1.5 |
| REMS latency (min) | 120.3±8.1 | 132.7±10.5 |
| SE | 94.6±0.5 | 93.5±1.3 |

TST, total sleep time; REMS, REM sleep; SE, sleep efficiency defined as TST/Time in Bed; WASO, wake after sleep onset

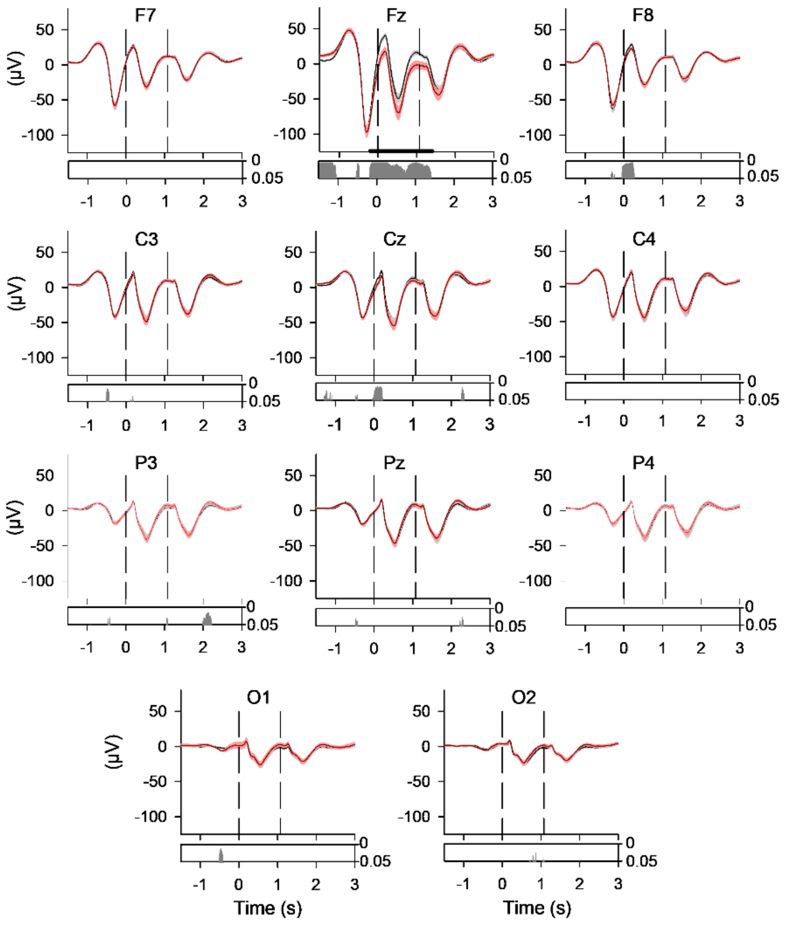


**Supplementary Figure 1**
Stimulus-locked EEG responses to acoustic stimuli in CLAS and CmodCLAS for all time periods of CLAS. Top diagrams: Grand mean waveforms (± SEM) at locations F7, Fz, F8, C3, Cz, C4, P3, Pz and P4 for CLAS (black) and CmodCLAS (red); baseline normalized data. Bottom diagrams: horizontal bars indicate time points (clusters) of significant differences (permutation test). N = 23? Baseline normalization occurred over -0.99 to -0.01 sec.

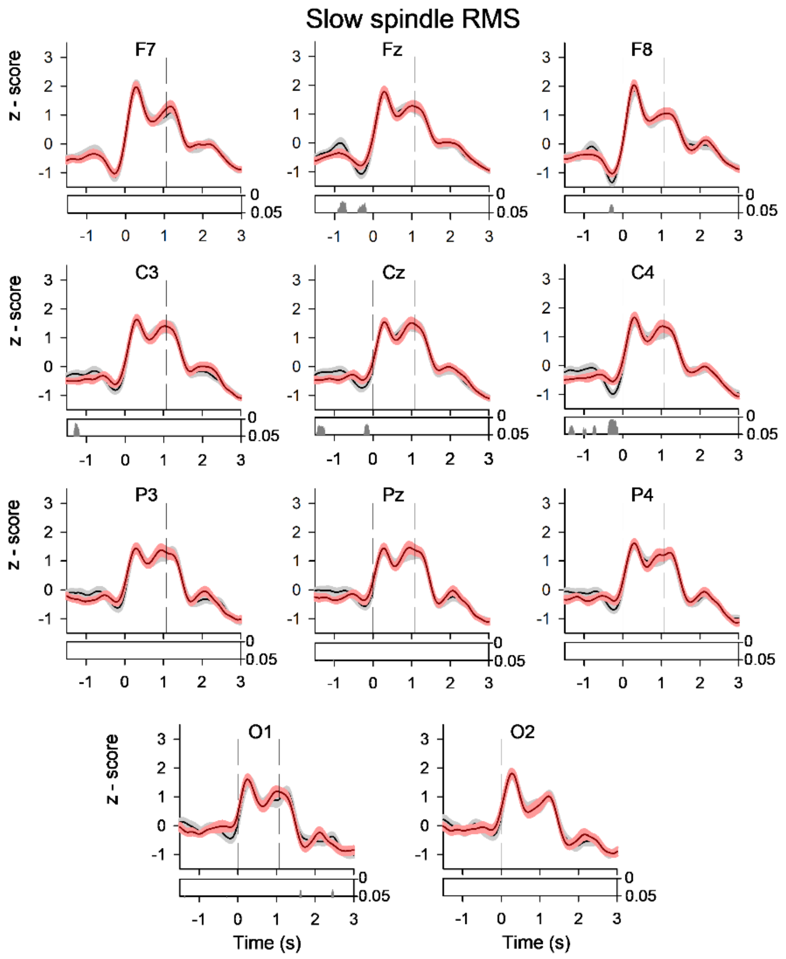


**Supplementary Figure 2**

Stimulus-locked slow spindle root mean square (RMS) to acoustic stimuli in CLAS and CmodCLAS.
Top diagrams: Grand mean waveforms (± SEM) of stimulus-locked responses at all 11 topographies for CLAS (black) and CmodCLAS (red). Bottom diagrams: horizontal bars indicate time points of significant differences without multiple comparison correction (Wilcoxon signed-rank test), N =23.

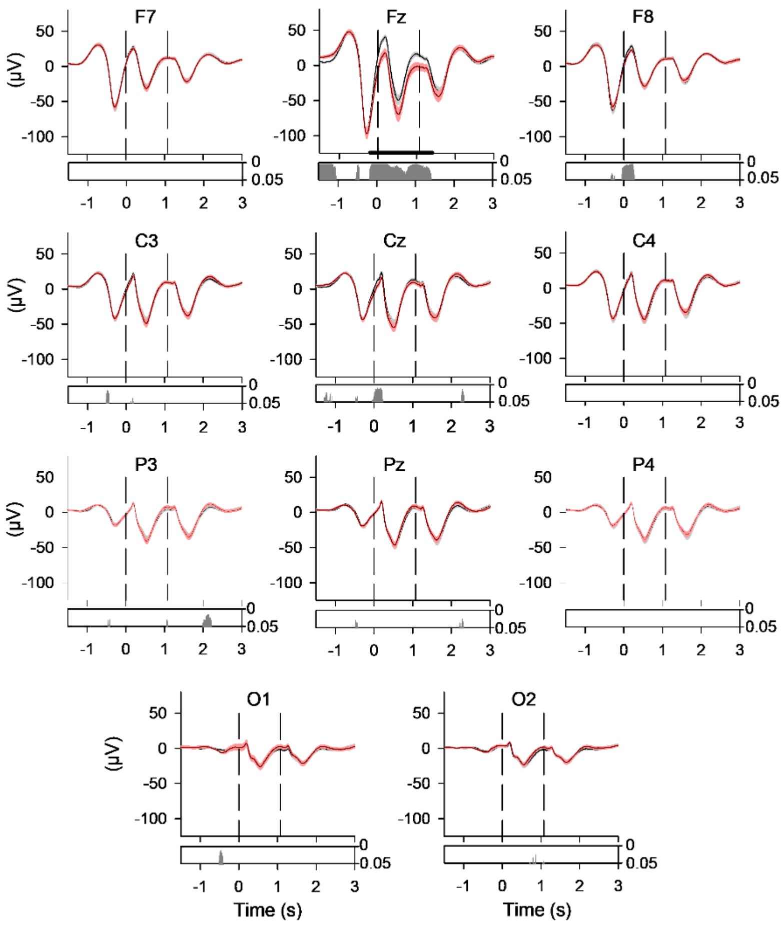


**Supplementary Figure 3**

Stimulus-locked fast spindle root mean square (RMS) to acoustic stimuli in CLAS and CmodCLAS.
Top diagrams: Grand mean waveforms (± SEM) of stimulus-locked responses at all 11 topographies for CLAS (black) and CmodCLAS (red). Bottom diagrams: horizontal bars indicate time points of significant differences (Wilcoxon signed-rank test, $p < 0.05$, uncorrected), N =23.

**Supplementary Table S2.** Slow and fast spindle properties during N2 and N3.

| Sleep Oscillation Properties | |
|---|---|
| Slow Spindle Density (per 30s; Fz) | 1.6±0.0 |
| Fast Spindle Density (per 30s; Cz) | 1.9±0.1 |
| Slow Spindle Length (s; Fz) | 991.8±13.3 |
| Fast Spindle Length (s; Cz) | 942.8±14.0 |
| Slow Spindle RMS (Fz) | 12.4±0.7 |
| Fast Spindle RMS (Cz) | 11.9±0.6 |

Data are averaged across CLAS and CmodCLAS for Fz and Cz, respectively.

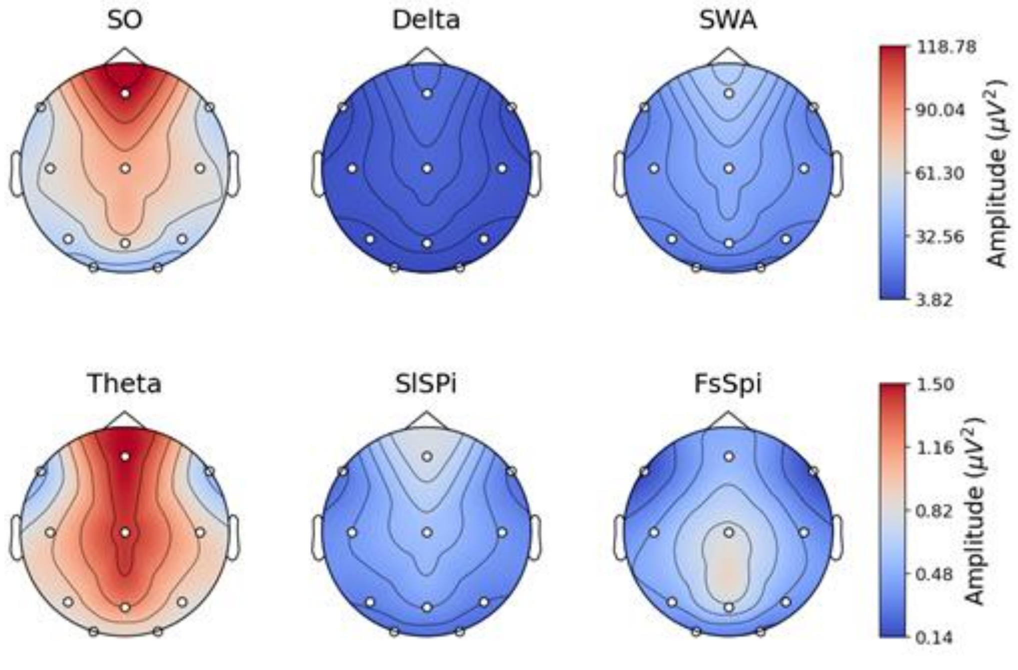


**Supplementary Figure 4**
Heat maps of EEG power, averaged across CLAS and CmodCLAS for the frequency bands of slow oscillation (SO), Delta, slow wave activity (SWA), theta, slow spindles (SlSpi), and fast spindles (FsSpi).

**Supplementary Table S3 Psychometric control tests**

| | **CLAS** | | **CmodCLAS** | |
|---|---|---|---|---|
| | Evening | Morning | Evening | Morning |
| SSS | 4.6±0.3 | 2.7±0.2 | 4.2±0.2 | 2.6±0.2 |
| PANAS-P | 2.1±0.1 | 2.7±0.1 | 2.4±0.1 | 2.8±0.1 |
| PANAS-N | 1.2±0.1 | 1.2±0.1 | 1.2±0.1 | 1.2±0.1 |

SSS, Stanford Sleepiness Scale; PANAS, Positive and Negative Affect scale
